# Mushroom body-specific profiling of gene expression identifies regulators of long-term memory in *Drosophila*

**DOI:** 10.1101/235960

**Authors:** Yves F Widmer, Adem Bilican, Rémy Bruggmann, Simon G Sprecher

## Abstract

Memory formation is achieved by genetically tightly controlled molecular pathways that result in a change of synaptic strength and synapse organization. While for short-term memory traces rapidly acting biochemical pathways are in place, the formation of long-lasting memories requires changes in the transcriptional program of a cell. Although many genes involved in learning and memory formation have been identified, little is known about the genetic mechanisms required for changing the transcriptional program during different phases of long-term memory formation. With *Drosophila melanogaster* as a model system we profiled transcriptomic changes in the mushroom body, a memory center in the fly brain, at distinct time intervals during long-term memory formation using the targeted DamID technique. We describe the gene expression profiles during these phases and tested 33 selected candidate genes for deficits in long-term memory formation using RNAi knockdown. We identified 10 genes that enhance or decrease memory when knocked-down in the mushroom body. For *vajk-1* and *hacd1*, the two strongest hits, we gained further support for their crucial role in learning and forgetting. These findings show that profiling gene expression changes in specific cell-types harboring memory traces provides a powerful entry point to identify new genes involved in learning and memory. The presented transcriptomic data may further be used as resource to study genes acting at different memory phases.

## INTRODUCTION

One of the most intriguing function of the brain is its ability to form long-term memories (LTMs), which may be stored for days, months or even a lifetime. The molecular and genetic process underlying LTM formation include de novo protein synthesis and corresponding changes in gene regulation, which in turn result in long-lasting changes in synaptic plasticity (Davis and Squire 1984; Tully *et al.* 1994). Initial studies in the Californian sea hare *Aplysia californica* identified the transcription factor cAMP response element binding protein (CREB), which is required for gene regulation of LTM formation (Dash *et al.* 1990; Lee *et al.* 2012). The importance of CREB in LTM formation has been confirmed in vertebrates and invertebrates, indicating its conservation through evolution, supporting that similar pathways control LTM formation (Yin and Tully 1996; Silva *et al.* 1998; Kandel *et al.* 2014). CREB is the most prominent example, but other transcription factors contribute to the regulation of transcription significant for memory and synaptic plasticity (Alberini 2009). New protein synthesis is not only required for LTM formation, but also at later phases after learning. Several studies have shown that reactivated or recalled memories become sensitive to disruption and that stabilization is again dependent on protein synthesis (Nader *et al.* 2000; Kida *et al.* 2002; Pedreira *et al.* 2002; Lee *et al.* 2012). In addition, a later wave of mRNA and protein synthesis seems to be essential for LTM maintenance (Bekinschtein *et al.* 2007; Katche *et al.* 2010). A recent study in *Drosophila* showed that CREB dependent transcription is also required for LTM maintenance, however a different coactivator interacts with CREB in memory formation and maintenance (Hirano *et al.* 2016). Moreover, late memory maintenance becomes independent of CREB, but requires other transcription factors. Although many genes involved in the acquisition and consolidation of memories have been identified, little is known about the genetic bases of long-term memory formation and maintenance. In *Drosophila*, most studies on LTM focused on the first 24 h time window after learning. Therefore, our understanding of the genetic and molecular mechanisms of LTM is mostly limited to this early time window.

The mushroom body (MB) represents the main center of olfactory associative memory in the *Drosophila* brain (Heisenberg *et al.* 1985; de Belle and Heisenberg 1994; Dubnau *et al.* 2001). Each MB contains about 2500 neurons, called Kenyon cells (KCs), that receive input from olfactory projection neurons and extend axons to form lobe structures (Aso *et al.* 2009). KCs are classified into three classes, α/β, α’/β’ and γ, according to their projection pattern in the lobes (Crittenden *et al.* 1998). Dopaminergic neurons from the protocerebral anterior medial (PAM) cluster convey the sugar reward signal to the MB, where the association of the odor and the reward is taking place (Liu *et al.* 2012; Burke *et al.* 2012). We here use a MB-specific transcriptomic approach to identify genes that are involved in long-term memory maintenance and forgetting. We made use of the Targeted DamID (TaDa) technique to profile transcription in KCs (Southall *et al.* 2013). TaDa enables cell-type specific gene expression profiling with temporal control. The system employs DNA adenine methyltransferase (Dam) from *E. coli*, which is fused to RNA polymerase II (Pol II). Expression of the fusion protein results in methylation of adenine in the sequence GATC in loci that are bound by Pol II, providing a readout of transcriptional activity. Methylated fragments can specifically be amplified and then sequenced. We prepared and sequenced samples of paired and unpaired trained flies at four time intervals, each with four biological replicates, to analyze gene expression changes within KCs. Differentially expressed genes of these four time intervals after conditioning were determined and 33 candidate genes were selected and tested in a long-term memory RNAi experiment. 10 RNAi lines that showed a lower or higher 48 h memory performance than the control line were identified. Two genes, *vajk-1* (*CG16886*) and *hacd1* (*CG6746*), were examined in more detail. Knockdown of *hacd1* in the MB resulted in enhanced LTM, however short-term or middle-term memory was not affected. *vajk-1* knockdown in the MB showed impaired memory at all tested memory phases in knockdown experiments and could be involved in memory formation.

## MATERIALS AND METHODS

### Fly strains

*Drosophila melanogaster* flies were reared on cornmeal medium supplemented with fructose, molasses and yeast. If not mentioned differently, flies were kept at 25° and exposed to a 12 h light – 12 h dark cycle. For the experiments with *tubGal80^ts^*, flies were raised at 18° and moved to 29° five days before conditioning.

Canton-S was used as wild-type (courtesy of R. Stocker). *UAS-Dam* and *UAS-Dam-Pol II* were obtained from Tony D Southall (Imperial College London) and *mb247-Gal4* was obtained from Dennis Pauls (University of Würzburg). Used *UAS-RNAi* lines were received from VDRC stock center (Dietzl *et al.* 2007) or Transgenic RNAi Project (TRiP) collection (Perkins *et al.* 2015) (see table S2 and S3 for stock numbers). *UAS-Dcr-2* (24644) and *tubGal80^ts^* (7019) were obtained from Bloomington stock center.

### Olfactory appetitive conditioning

The memory apparatus used to conduct the behavior experiments is based on Tully and Quinn (1985) and was modified to allow performing four experiments in parallel. Two odors were used; limonene (Sigma-Aldrich, 183164) and benzaldehyde (Fluka, 12010). 85 μl of limonene was filled in plastic containers measuring 7 mm in diameter and 60 μl of benzaldehyde was filled in plastic containers measuring 5 mm in diameter. A vacuum pump adjusted to a flow rate of 7 l/min was used for odor delivery. Experiments were done at 22-25° and 70-75% relative humidity. Training was performed in dim red light and tests were performed in darkness. Filter paper soaked with a 1.5 M sucrose (Sigma-Aldrich, 84100) solution or distilled water were prepared the day before the olfactory conditioning experiments and left to dry at room temperature. 19-21 h before conditioning, groups of 60-100 flies (1-4 d old) were put in starvation vials and kept at 18°. Empty fly vials with wet cotton wool on the bottom were used to starve the flies.

For appetitive conditioning, starved flies were loaded in tubes lined with water filter papers. After an initial phase of 90 s, one of the odors was presented for 120 s. Then, flies were exposed for 60 s to non-odorized airflow. During this 60 s, flies were transferred to tubes lined with sucrose filter papers. Afterwards, the second odor was presented for 120s. To assess 0 h memory, flies were tested immediately after conditioning. For 3 h and 48 h memory, flies were put back in starvation vials and were kept at 18° until the test. Flies tested 48 hours after conditioning were put 21-23 h before the test on food for two hours. One experiment consisted of two reciprocal conditionings, in which the odor paired with sucrose was exchanged.

For the unpaired training protocol, flies were loaded in tubes lined with sucrose filter papers and after 120 s transferred to tubes lined with water filter papers. Two minutes after the sucrose, flies were exposed for 120 s to the first odor, 60 s to non-odorized airflow and 120 s to the second odor.

### Memory tests

For the memory test, flies were loaded into a sliding compartment and moved to a two-arm choice point where they could choose between benzaldehyde and limonene. After 120 s, gates were closed and the number of flies within each arm was counted. A preference index was calculated as follows:

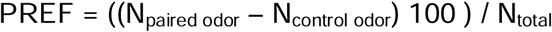

The preference indices from the two reciprocal groups were averaged to calculate a memory performance index (PI).

### Sensory tests

Sensory tests were performed with the same apparatus as the memory tests. To measure sucrose response, flies could choose between a tube lined with a sucrose filter paper and a tube lined with a water filter paper for 120 seconds. A sucrose preference index (PrefI) was calculated with this formula:

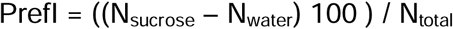

For the odor avoidance tests, flies could choose between a tube with an odor container attached (filled with benzaldehyde or limonene) and a tube with an empty plastic container attached. Flies in each tube were counted after 120 s and an odor preference index was calculated:

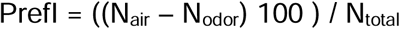

### Targeted DamID

*UAS-Dam* and *UAS-Dam-Pol II* flies were crossed to *tubGal80^ts^; mb247-Gal4.* Offspring of those crosses were reared at 18° and trained with the standard (paired) or unpaired olfactory appetitive conditioning paradigm. Expression of Dam and Dam-Pol II was induced by shifting the flies to 29°. Four time intervals were used: T1-T4. For T1, flies were moved to 29° three hours before conditioning, were trained at 25° and moved back to 29° for 12 h. For T2-T4, animals were trained at 18° and shifted to 29° 12-24 h (T2), 24-48 h (T3) or 48-72 h (T4) after conditioning. At the end of the 29°-time interval, flies were frozen in liquid nitrogen and heads were collected. Extraction of genomic DNA from the fly heads (50-100 per sample), amplification of methylated fragments, DNA purification and sonication was performed according to Marshall *et al.* (2016). After sonication, DamID adaptors were removed by digesting overnight at 37° with five units of Sau3AI (NEB, R0169S). The sequencing libraries were prepared according to the Illumina TruSeq nano DNA library protocol. The samples were sequenced using NGS (Illumina HiSeq3000) at an average of around 30 million paired-end reads per sample.

### Targeted DamID analysis

Low-quality bases of the sequencing reads were trimmed using Trimmomatic (Bolger *et al.* 2014) and the remaining good quality reads were mapped to the *D. melanogaster* genome (Release 6.05, Hoskins *et al.* 2015) using Bowtie2 (Langmead and Salzberg 2012). Next, the damidseq_pipeline software was used to process the data (Marshall and Brand 2015) and to generate log2 ratio files for each pairwise comparison between Dam-Pol II and Dam-only for each time point separately. For each gene the log2 ratio was calculated and a false discovery rate (FDR) assigned. The FDR value was generated via 50 000 simulations and represents the probability of having a given expression for a given gene based on the length of the gene and the total number of GATC sites present in this gene. Genes with a positive log2 ratio and a significant FDR value (FDR<0.05) were defined as expressed. Median log2 fold change values were compared and a Student’s *t*-test performed to identify differentially regulated genes between the paired and unpaired trained groups.

### Statistics

To compare PrefIs or PIs of two groups the Welch two sample *t*-test was used. To test if PI mean values are different from zero, the one sample Student’s *t*-test was used. Statistical analyses and graphical representation of the data were performed using R version 3.4.1 (R Foundation for Statistical Computing).

## RESULTS

### Assessing gene expression profiles during long-term memory formation

To study temporal gene expression changes in the mushroom body during long-term memory formation we used the Targeted DamID technique, an adaptation of the DNA adenine methyltransferase identification (DamID) technique (Steensel and Henikoff 2000; Southall *et al.* 2013). This technique employs an *Escherichia coli* DNA adenine methyltransferase (Dam), which is fused to a DNA-associated protein of interest. The bacterial Dam methylates adenine in the GATC sequence, tagging the regions of the genome where the Dam-fusion protein has interacted with DNA. TaDa allows temporally controlled expression of Dam in a cell- or tissue-specific fashion. We used TaDa to profile RNA polymerase II binding, which allows the identification of actively transcribed loci and therefore provides an indirect readout of gene expression. The Dam-Pol II fusion protein (*UAS-Dam-Pol II*) was expressed in Kenyon cells of the mushroom body, a brain centre for olfactory associative memory, using the *mb247-Gal4 (MB-Gal4)* driver (Heisenberg *et al.* 1985; de Belle and Heisenberg 1994; Dubnau *et al.* 2001). *mb247-Gal4* drives expression of UAS-target transgenes in the α, β and γ lobes of the mushroom body (Zars *et al.* 2000; Aso *et al.* 2009). In order to control for unspecific methylation we compared the expression of Dam-Pol II with UAS-Dam. Temporal restricted expression was achieved using the temperature sensitive *tubulin-Gal80^ts^.*

To study long-term memory, flies were trained in a classical olfactory conditioning paradigm, using sucrose as a positive reinforcer. Animals were sequentially exposed to two odorants, one of which was paired with sucrose. Following the learning procedure, flies preferentially moved towards the paired odor. A single trail of appetitive olfactory conditioning is capable of inducing LTM that lasts for days (Krashes and Waddell 2008; Colomb *et al.* 2009). In an unpaired training protocol, in which odors and sucrose were presented temporally separated, flies did not form odor memories (Figure 1A and B). To gain insight into the transcriptional changes underlying the formation, consolidation and maintenance of LTM, we performed a TaDa sequencing analysis during different time intervals after training. Gene expression was compared between flies that were trained to associate sucrose with an odor and control flies that received sucrose and odors unpaired in time. The experiment was designed as a time course with four time intervals after the conditioning protocol (Figure 1D). Time interval 1 (T1) included the first 12 hours after training. Flies were moved to 29° three hours before olfactory conditioning to induce expression of Dam-Pol II or Dam-only in the MB. The second time interval (T2) was 12 to 24 h, T3 was 24 to 48 h and T4 was 48 to 72 h after training. For each time point we used four biological replicates for experiment and control (Dam-Pol II and Dam-only) as well as for paired and unpaired training. At the end of the induction time interval the heads of the flies were collected. Genomic DNA was extracted from heads and digested with the restriction enzyme Dpn I, which cuts at adenine-methylated GATC sites. Methylated fragments were PCR amplified and DNA was prepared and sequenced using Illumina HiSeq3000 with about 30 million paired-end reads per sample in average (Figure 1C). Next, we calculated the log2 ratio between Dam-Pol II and Dam-only samples. A positive log2 ratio of Dam-Pol II /Dam-only implies that GATC sites were preferentially methylated in Dam-Pol II samples compared to Dam-only background methylation, indicating active transcription of a given gene locus (Figure 1C). We further calculated for each gene from a given condition at a given time interval expression values and false discovery rates (FDRs). The FDR value represents the probability of having a given expression for a given gene based on the length of the gene and the total number of GATC sites present in this gene. Based on the expression level and the FDR value, we identified genes as expressed in case of a positive log2 ratio and a significant FDR value (FDR<0.05). To find differentially expressed genes between paired and unpaired trained groups, calculated expression values were compared and p-values determined with a Student’s *t*-test. At T1 we identified 86 differentially expressed genes. Of those, 46 were upregulated and 40 were downregulated in the paired group compared to the unpaired group (Figure 1E). For T2, 115 genes were significantly higher expressed in the paired trained group. 56 differentially upregulated genes and 45 downregulated genes were found at T3. At the last time interval (T4), we identified 75 genes with higher and 202 genes with lower median expression in the paired conditioned group (Table S1).

**Figure 1.**
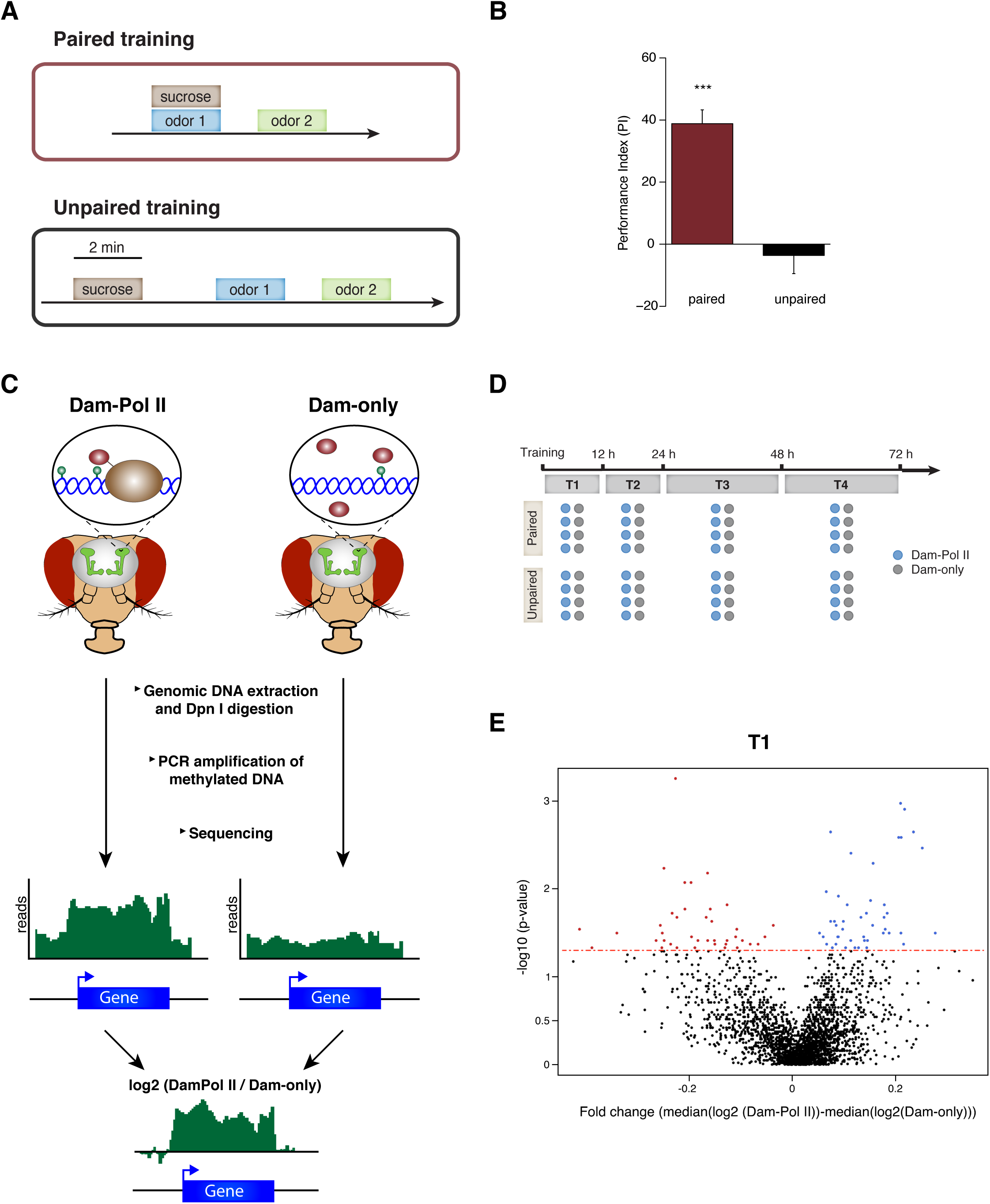
Monitoring gene expression changes after memory formation. (A) Illustration showing the two used learning paradigms. For the paired training, flies were allowed to feed on sucrose during odor 1 presentation. For the unpaired training, the sucrose feeding was separated from the odor presentations. (B) Paired trained wild-type flies displayed long-term memory measured after 24 h. Flies exposed to the unpaired training did not show a changed odor preference. Bar graphes represent the mean and error bars represent the standard error of the mean (SEM). N=8. ***p<0.001 (one-sample *t*-test). (C) Schematic representation of the targeted DamID (TaDa) experimental pipeline. A mushroom body specific Gal4 driver line was used to drive expression of *Dam-Pol II* and *Dam-only.* Genomic DNA from heads was extracted and digested with the methylation sensitive restriction enzyme DpnI. Methylated sequences were then amplified by PCR and sequenced. The resulting reads were mapped to a reference genome and the log2 ratio of Dam-Pol II/Dam-only was calculated. A positive ratio indicates transcription by the RNA Polymerase II. (D) Schematic illustration of the experimental design to profile gene expression after memory formation. Transcription was monitored at four time intervals (T1-T4) after training. Four replicates for *Dam-Pol II* and *Dam-only* expressing flies were conducted at each time interval with flies trained in a paired and flies trained in an unpaired conditioning paradigm. (E) Representative volcano plot showing differentially regulated genes between paired and unpaired trained flies at the first time interval (T1). Median log2 fold changes are plotted against –log10 (p-value). Genes with a significant fold change between paired and unpaired are colored. Red indicates significant higher expression in the unpaired group and blue indicates significant higher expression in the paired group. The dashed red line indicates a p-value of 0.05.

### Candidate RNAi screen for long-term memory defects

To identify genes that regulate long-term memory formation, maintenance and forgetting we screened candidate genes for LTM defects using MB-specific *UAS-RNAi*. We selected a total of 33 candidate genes based on their expression profile in the MB during long-term memory formation. Selected candidate genes could be positive regulators that form or stabilize memories as well as negative regulators that hinder formation and maintenance or actively remove memories. We therefore generated a ranked list of differentially up- and downregulated genes for different time intervals (Table S1). The ranking was based on the median gene expression differences between the paired and unpaired group. For T3 and T4 the six highest ranked genes as upregulated and downregulated were selected. Since no significant downregulated genes were identified for T2 we selected the top ten ranked upregulated genes at T2. We further selected eight genes according to gene ontology (GO) terms (Ashburner *et al.* 2000; The Gene Ontology Consortium 2017): transcription factor activity (*E*(*spl*)*mβ*-*HLH*, *Cdk7* and *Hsf*), actin cytoskeleton organization (*Vps4* and *capt*), cellular component of dendrites (*Mmp1*); synaptic vesicle docking (*Syx8*) and a gene implicated in LTM (*Hn*).

In 2015, Walkinshaw *et al.* performed a large genetic screen analysis, testing 3200 RNAi lines for 3 h memory, in which 42 genes that enhance memory were identified. Three of these genes (*amon, prt* and *hacd1*) were differentially expressed at T2, T3 or T4 in our experiment, which we added to our selection of candidate genes. Those three genes could possibly also be negative regulators of LTM.

Flies expressing *UAS-RNAi* under the control of *mb247-Gal4* were trained using the same appetitive olfactory learning paradigm as above for TaDa experiment and LTM performance was assessed 48 hours later. We chose to test for 48 h memory to identify genes that are involved in LTM. To increase efficiency of RNAi we coexpressed a *UAS-Dcr2* transgene (Dietzl *et al.* 2007). The microRNA (miRNA) mir- 282 was inhibited with a miRNA sponge construct (Fulga *et al.* 2015).

As positive control, we included a *UAS-RNAi* knockdown against the adenylyl cyclase *rutabaga (rut)*, which is required for memory formation (Blum *et al.* 2009). We indeed observed that a MB-specific knock-down of *rut* resulted in impaired memory performance (Figure 2A). Three *mb247-Gal4/UAS-RNAi* crosses did not produce viable adult offspring, thus could not be tested for memory performance. The offspring of the remaining lines did not display visible developmental defects. Tested *UAS-RNAi* lines with +/− one standard deviation from the performance index (PI) of the driver line *MB-Gal4* were selected as positive hits. 20 RNAi lines showed no significant changes in memory performance, while one RNAi line showed a decreased memory performance and nine RNAi lines an increased 48 h memory performance (Figure 2A; Table S2). None of these genes were previously reported to be involved in LTM and therefore represented new genes potentially regulating LTM. To further support the function of these 10 genes in LTM we re-tested them using a second different *UAS-RNAi* line. No second RNAi line against *Cpr64Aa* was available and therefore we did not test this gene in the retest experiment. For four genes we could again observe a memory performance different from *MB-Gal4* (p-value<0.05, Welch two sample *t*-test). Knockdown of the gene *vajk-1 (CG16886)* resulted again in reduced LTM and knockdown of *CG12338, hacd1 (CG6746)* and *CG14572* caused memory enhancement. Four RNAi lines performed not significantly different from the driver line and for two genes no second construct was available (Figure 2B, Table S3).

**Figure 2.**
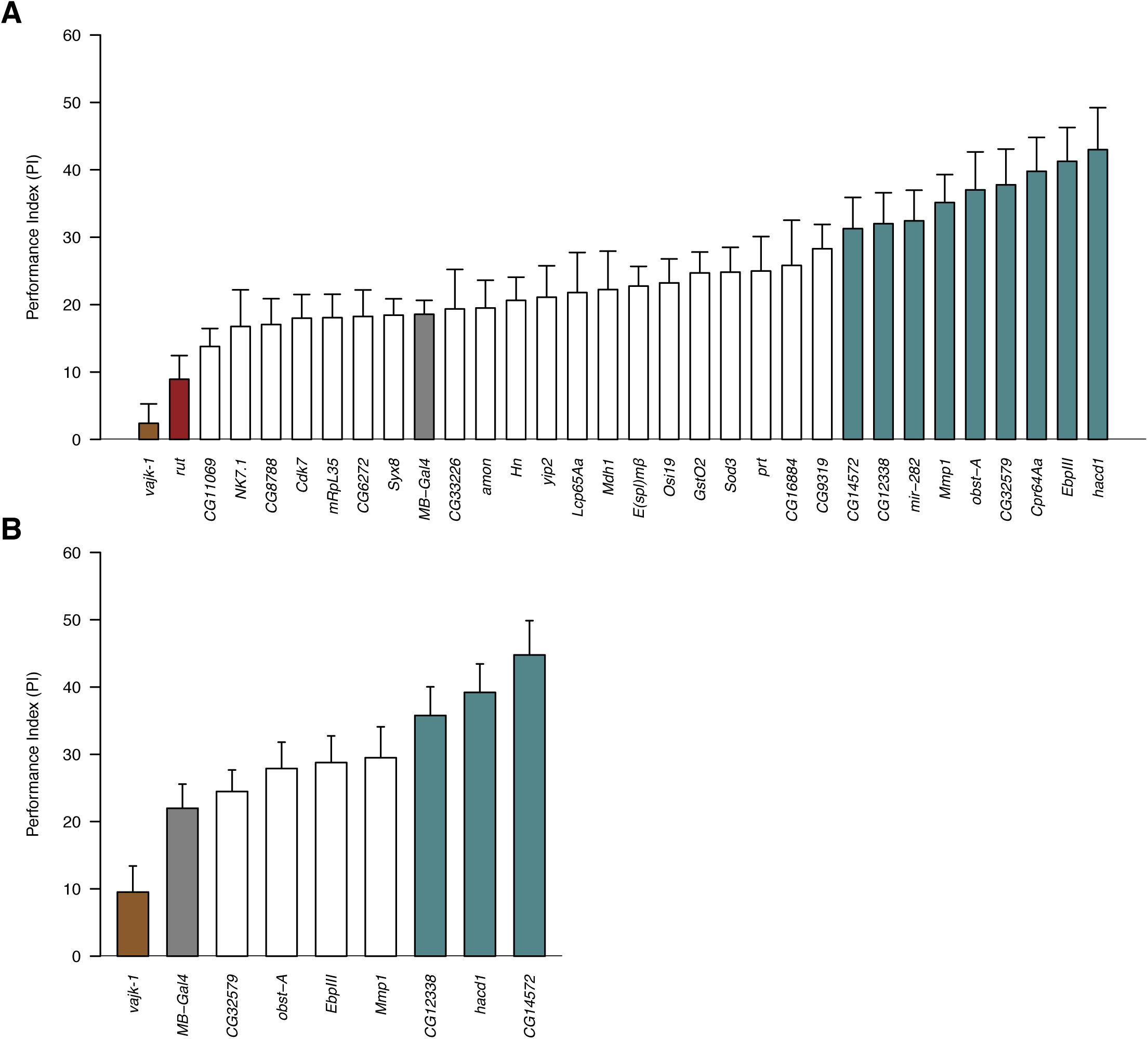
48 h memory RNAi screen. Flies were trained in an appetitive olfactory conditioning paradigm and memory was assessed after 48 h. RNAi constructs were used to inhibit gene products of the candidate genes. (A) RNAi lines with at least +/- one standard deviation from the performance index (PI) of the driver line *MB-Gal4* (highlighted in gray) were defined as hits. Nine RNAi lines with a higher memory performance (highlighted in teal blue) and one RNAi line with a lower memory performance (highlighted in brown) were identified. *rutabaga (rut)* was added as a positive control (highlighted in red). N=6-8 for RNAi lines, N=20 for *MB-Gal4.* (B) Hits were reevaluated using another RNAi line targeting those genes. RNAi lines, which performed significantly different from the driver line *MB-Gal4,* are depicted in brown and teal blue. N=8-9. Bar graphs represent the mean and error bars represent the standard error of the mean (SEM).

### Knockdown of Hacd1 in the mushroom body increases long-term memory

We next further assessed the top hit with increased or decreased 48 h memory in more detail. We first analyzed *hacd1*, which encodes 3-hydroxyacyl-CoA dehydratase (HACD) an enzyme in lipid metabolism, required for catalyzing very long-chain fatty acid (VLCFA) (Denic and Weissman 2007; Ikeda *et al.* 2008). A previous study provides genetic evidence for *hacd1* in VLCFA biosynthesis, since RNAi knockdown of *hacd1* in oenocytes led to a strong decrease in cuticular hydrocarbon levels (Wicker-Thomas *et al.* 2015). In the initial MB-specific RNAi screen *hacd1* showed the highest LTM score, which was confirmed by a second *UAS-RNAi* line (Figure 2). To further validate this finding we tested *mb247-Gal4/UAS-hacd1-RNAi* alongside with both parental control strains. Moreover, we wondered if this enhanced memory is specific to LTM or also manifests itself in earlier memory phases. Thus, in addition to long-term memory (48 h) we tested middle-term memory (3 h) and short-term memory, measured directly after training (0 h). No significant differences between the groups were detected for 0 h and 3 h memory, indicating that short-term and middle-term memory are not affected by *hacd1-RNAi* knockdown (Figure 3). However, we observed again a significant higher PI in *mb247-Gal4/UAS-hacd1-RNAi* flies compared to control parental lines, supporting a role of *hacd1* in LTM.

**Figure 3.**
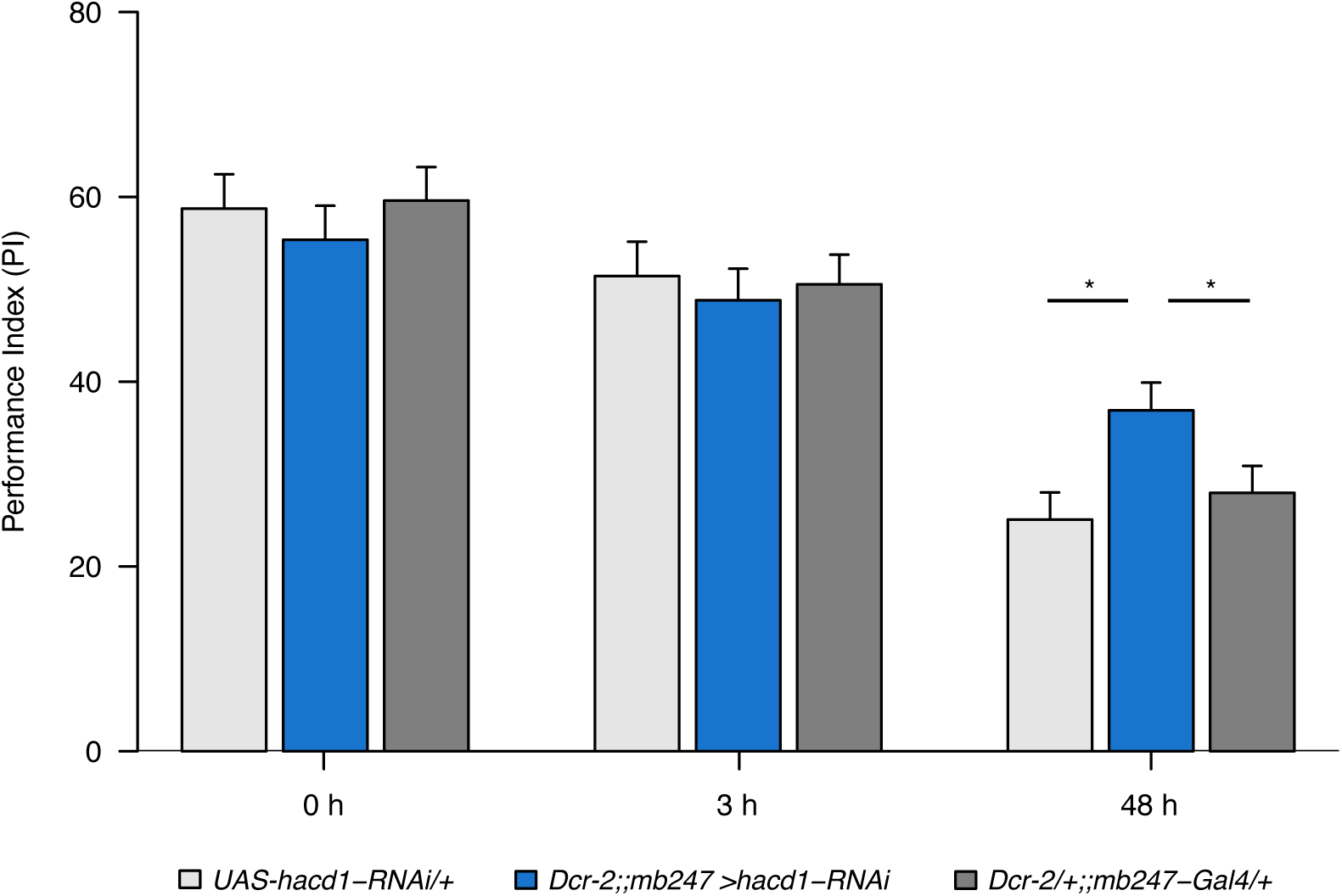
Knockdown of *hacd1* in the mushroom body enhances 48 h memory. Flies expressing *hacd1-RNAi* in Kenyon cells were tested along with controls for their memory performance immediately after conditioning (0 h) or after a period of 3 h and 48 h. Memory measured two days after training was significantly enhanced in *hacd1-RNAi* expressing flies compared to parental lines. Performance indices (PIs) did not differ between groups after 0 h and 3 h. N=9-12. Bar graphs represent the mean and error bars represent the standard error of the mean (SEM). Asterisks denote significant difference between groups (* p<0.05).

### Knockdown of vajk-1 in the mushroom body results in learning defects

The *vajk-1* gene is located within the Nimrod gene cluster on chromosome 2, the largest syntenic unit in the genome (Somogyi *et al.* 2010; Cinege *et al.* 2017). Genes of the Nimrod cluster have been suggested to be contributing to innate immune response (Kurucz *et al.* 2007). MB-specific expression of *UAS-vajk-1-RNAi* resulted in a low 48 h memory performance, which was confirmed by a second *UAS-RNAi* line (Figure 2). We next assessed if *vajk-1* knockdown affects only LTM by testing short-term and middle-term memory. We found that *mb247-Gal4/UAS-vajk-1-RNAi* flies showed impaired short-, middle- and long-term memory by displaying significantly lower memory scores at all three time points, when compared to parental control strains (Figure 4A). The finding that *UAS-vajk-1-RNAi* expressing flies showed reduced memory immediately after learning, suggests that *vajk-1* may be involved in memory formation. Reduced learning capability may also result from developmental defects in MB formation or defects in sensory input. We therefore performed sensory tests with *mb247-Gal4/UAS-vajk-1-RNAi* flies and the parental lines. All the tested lines moved away from the presented odors (benzaldehyde or limonene) and no significant difference between the groups was observed (Figure 4B,C). *UAS-vajk-1-RNAi* expression also had no influence on the sugar attraction behavior. Sucrose response was not different from the control lines (Figure 4D). To examine if developmental defects cause the observed memory impairment, we temporally restricted RNAi expression with *tubGal80^ts^.* Flies were raised at 18° and moved after hatching to 29° for five days to induce expression of the RNAi construct. Memory was assessed 0 h and 48 h after memory formation. Knockdown of *vajk-1* in the MB induced reduced learning, measured directly after conditioning. For 48 h memory, flies expressing *UAS-vajk-1-RNAi* showed a lower performance than the parental Gal4 control, but the memory score was not significantly different from the parental UAS control (p-value=0.18, Welch two sample *t*-test) (Figure 4E).

**Figure 4.**
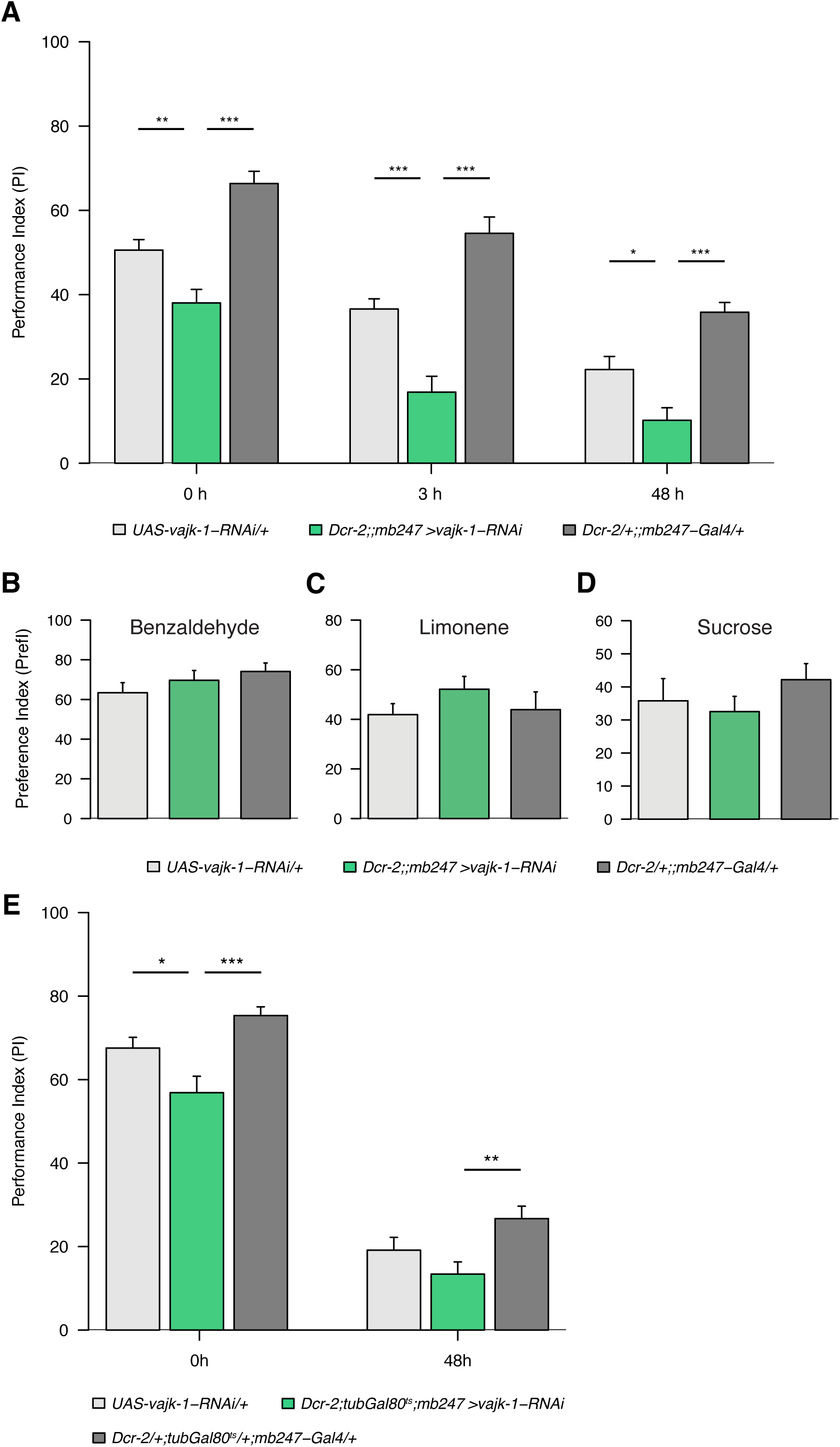
Expression of *vajk-1-RNAi* in Kenyon cells causes memory impairment. (A) Flies were trained in an appetitive olfactory learning paradigm and tested 0 h, 3 h or 48 h later. Inhibiting *vajk-1* in the mushroom body resulted in reduced memory performance at all measured time points compared to parental lines. N=9-13. (B, C) Benzaldehyde and limonene odor avoidance of the parental lines and the *vajk-1-RNAi* line crossed to *mb247-Gal4* were tested. No significant differences between the groups were observed. N=8-9. (D) Flies with RNAi inhibited *vajk-1* performed not significantly different from the control lines in a sucrose response test. N=12-13. (E) A late knockdown of *vajk-1* was achieved by shifting the flies after hatching to 29°C to activate expression of *vajk-1-RNAi.* Knockdown of *vajk-1* resulted in reduced learning performance directly after training compared to parental lines. 48 h after training *vajk-1-RNAi* expressing flies displayed a lower memory performance index than the parental Gal4 control line, but did not significantly differ from the parental UAS control line (p-value=0.18). N=11-16. Bar graphs represent the mean and error bars represent the standard error of the mean (SEM). Asterisks denote significant difference between groups (* p<0.05, ** p<0.01, ***p<0.001).

## DISCUSSION

Appetitive olfactory learning using sugar as reward forms memories, which last for several days after training. To monitor the transcriptional program that occurs in the mushroom body we used a transcriptomics approach identifying actively transcribed genetic loci during four time intervals after training. Based on the analysis of MB gene expression profiles we performed a MB-specific candidate RNAi screen, which identified 10 genes that exhibited altered 48 h memory performance in an RNAi knockdown experiment, confirming four in a re-screen using independent *UAS-RNAi* lines. The two genes *vajk-1* and *hacd1*, top candidates of increased and decreased 48 h memory, were further tested in more detail.

### Transcriptomics characterization of gene expression profiles during LTM in the MB

Genetic tools available in *Drosophila* provide a fruitful basis to study genetic mechanisms required for different phases of learning and memory formation (Keene and Waddell 2007). We here used the TaDa technique, which enabled us to specifically profile gene expression in the MB, without the requirement of isolating cells from this brain structure. Our findings show that TaDa with Dam-Pol II is a valuable technique to measure gene expression in a specific cell population in the *Drosophila* brain. TaDa is a quite recent technique and has not been used before to study memory related gene expression changes. The bioinformatics analysis of sequencing data allowed us to identify changes in the transcriptome at the whole-genome level during different phases of memory formation between paired and unpaired trained flies. A functional RNAi screen supports the validity of this technical approach (see below). While we here specifically focused on the later stages during LTM formation and forgetting, the transcriptome data further allows studies on genetic aspects of memory initiation, thus providing a valuable resource for future functional studies.

### RNAi based identification of novel genes in regulating LTM forgetting

From the selected 33 candidate genes for the *UAS-RNAi* screen for LTM we identified one hit with lower and nine hits with higher 48 h memory performance, thus roughly 30% of the selected candidates were identified as hits. Since RNAi lines may identify false positive genes by off-target effects we further used a second *UAS-RNAi* line for eight of the hits, out of which four showed again a significantly altered LTM performance. Thus, we confirmed four out of 33 candidate genes as novel genes in memory formation, maintenance or forgetting. It is important to note that *UAS-RNAi* induced knockdown may reduce protein levels only partly and therefore cause a hypomorphic phenotype resulting in a false negative call. Since about 50% of the hits of the first RNAi screen did not show a phenotype in the re-screen, it seems likely that more candidate genes may actually cause a behavioral phenotype when stronger gene inactivation is used. In the future other techniques may be used to study these genes, including Minos transposon insertions to tag and knockdown proteins (Nagarkar-Jaiswal *et al.* 2015), the generation of transgenic CRISPR/Cas9 to generate mutations in the locus (Xu *et al.* 2015) or *UAS-ORF* lines for overexpression experiments (Bischof *et al.* 2013).

Interestingly, we found nine candidate genes with enhanced LTM performance. Reducing the effect of the microRNA mir-282, by using a miRNA sponge construct, showed increased 48 h memory when expressed in KCs (Fulga *et al.* 2015). Other miRNAs have previously been identified in learning and memory formation in *Drosophila.* While inhibiting mir-980 showed enhanced short-term and middle-term memory, mir-276a has been described to be necessary for LTM formation (Li *et al.* 2013; Guven-Ozkan *et al.* 2016). Interestingly while both genes regulated neuronal excitability, the molecular mechanisms that regulate learning and memory appear distinct. The autism susceptibility gene, *A2bp1* was identified as target of mir-980 causing memory enhancement, while mir-276a interferes with memory formation by regulating Dopamine receptor expression (Li *et al.* 2013; Guven-Ozkan *et al.* 2016). While the role for mir-282 in learning and memory formation was unknown, a recent study identified the adenylyl cyclase *rutabaga* as target gene of mir-282 (Vilmos *et al.* 2013). Moreover, using the microRNA.org resource for miRNA target prediction, we found that five of the top 25 target genes are reported to be involved in learning and memory (Betel *et al.* 2008, 2010) (Table S4).

*CG12338* encodes a protein, which was suggested to be involved in the D-amino acid metabolic process (Gaudet *et al.* 2011). The mouse homolog *D-amino acid oxidase (Dao)* has a critical role in spatial memory. Mutant mice lacking DAO performed significantly better than wild-type mice in the Morris water maze test (Maekawa *et al.* 2005). *Dao* mutant mice have a higher D-amino-acid concentration in the brain, which possibly enhances N-methyl-D-aspartate (NMDA) receptor response and thereby facilitates spatial learning (Hashimoto *et al.* 1993; Morikawa *et al.* 2001). Our observation that knocking-down *CG12338* caused enhanced memory suggests that similar mechanism may be involved in regulating memory in *Drosophila*.

Two of our memory enhancing hits regulate extracellular matrix organization: *obstructor-A* (*obst-A*) and *Matrix metalloproteinase 1* (*Mmp1*). Obst-A was shown to be required for ECM dynamics and coordination of ECM protection (Petkau *et al.* 2012). Mmp1 belongs to a conserved family of extracellular proteases that cleave protein components of the ECM. Mmp1 can mediate matrix remodeling and is required for degrading severed dendrites during metamorphosis (Kuo *et al.* 2005; Glasheen *et al.* 2010). In rats, it was observed that MMP-3 and −9 increased learning-dependent and inhibition altered long-term potentiation and learning capacity (Meighan *et al.* 2006). *Cuticular protein 64Aa (Cpr64Aa),* another gene that showed a higher 48 h memory performance in the RNAi screen, is also reported to be a cellular component of the ECM. The ECM is a dynamic structure that can alter the synaptic efficiency, thus contributing to synaptic plasticity (Wlodarczyk *et al.* 2011; Frischknecht and Gundelfinger 2012). Specialized structures of stable and accumulated ECM molecules called perineuronal nets (PNNs) were found around certain neurons in the mammalian brain where they play a critical role in control of plasticity (Härtig *et al.* 1992; Pizzorusso *et al.* 2002). PNNs were shown to participate in memory mechanisms and modifications of PNNs can enhance long term memory (Gogolla *et al.* 2009; Romberg *et al.* 2013; Hylin *et al.* 2013). Digestion of PNNs mediated prolonged long-term object recognition memory and the same prolongation was observed in mice lacking an essential PNNs component (Romberg *et al.* 2013). It has been suggested that long-term memories could be stored and maintained in neuron surrounding ECM structures (Tsien 2013). Our results suggest that in *Drosophila* ECM proteins could also contribute to memory maintenance. The discovered genes will serve as a valuable starting point for future studies of the molecular mechanisms underlying LTM.

### Identification of *vajk-1* and *hacd1* as learning and memory genes

Expression of *UAS-vajk-1-RNAi* in the MB caused memory impairment. The *vajk-1* gene is located together with two homologous genes (*vajk-2* and *vajk-3*) in a large intron of *Ance-3,* which is part of the Nimrod cluster. However, *vajk* genes are not related to the Nimrod genes, which are involved in the innate immune defense. The *vajk* gene members are conserved in insects, but their function is unknown (Somogyi *et al.* 2010; Cinege *et al.* 2017). Our results suggest that *vajk-1* could be involved in the memory formation process.

We found that RNAi knockdown of *hacd1* in Kenyon cells resulted in enhanced 48 h memory, but 0 h and 3 h memory were unaffected. *hacd1* was one of 42 identified genes that showed increased 3 h memory after RNAi knockdown (Walkinshaw *et al.* 2015). Our results did not show memory enhancement 3 h, but 48 h after memory formation. This discrepancy is probably due to the use of a different driver line or learning paradigm. We used a mushroom body driver line and conditioned the flies in an appetitive paradigm. In contrast, Walkinshaw et al. (2015) used the panneuronal driver *Nsyb-Gal4* and aversive olfactory conditioning.

*hacd1* is involved in the synthesis of VLCFA and catalyzes the dehydration of the 3-hydroxyacyl-CoA (Wicker-Thomas *et al.* 2015). *hacd1* is evolutionary conserved among eukaryotes, but little is known about its function. Mammals have two homologs; *HACD1* and *HACD2.* Expression of *HACD2* was shown to be ubiquitous, whereas *HACD1* was found in heart and muscle cells and linked to certain muscle diseases and arrhythmogenic right ventricular dysplasia (Li *et al.* 2000; Wang *et al.* 2004; Pelé *et al.* 2005). The yeast homolog *PHS1* is also involved in the fatty acid elongation process, responsible for the third step in the VLFA synthesis cycle (Denic and Weissman 2007). Moreover it was proposed that *PHS1* is part of the endoplasmic reticulum membrane and possesses six transmembrane domains and (Kihara *et al.* 2008) and that it could be implicated in protein trafficking (Yu *et al.* 2006).

How *hacd1* could be implicated in LTM is currently unknown. However, various reports show that fatty acids and their mediators have numerous functions in the brain, including roles in learning and memory. It has been shown that overexpression of the fatty-acid binding protein *(Fabp)* in fruit flies increased LTM consolidation (Gerstner *et al.* 2011). Also in mammals, proteins involved in fatty acid metabolism can act on memory. It has been demonstrated that deletion of monoacylglycerol lipase caused memory enhancement in mice (Pan *et al.* 2011) and that inhibition of fatty acid amide hydrolase enhanced learning in rats (Mazzola *et al.* 2009).

Besides of lipid metabolism, the *Drosophila* Hacd1 protein could also be part of a signaling cascade, since it contains a protein-tyrosine phosphatase-like (PTPLA) domain that catalyzes the removal of a phosphate group attached to tyrosine. However, it has not been tested if this domain is functional. Future studies will be required to reveal the molecular mechanism of Hacd1 in LTM regulation.

## Data availability

Sequencing data can be accessed on BioProject (PRJNA419677).

## ACKNOWLEDGEMENTS

We would like to thank R. Stocker, T.D. Southall, D. Pauls, Bloomington stock center and VDRC stock center for fly strains. We also thank the Next Generation Sequencing Platform of the University of Bern for performing library preparation and sequencing experiments. We are also grateful to colleagues at the University of Fribourg and University of Bern for valuable discussions. This work was supported by SystemsX.ch (SynaptiX RTD) to SGS and RB.

---

**Table S1. Differentially regulated genes.**

Table showing differentially regulated genes between paired and unpaired trained flies. Genes are ranked according to the median gene expression difference.

**Table S2. Results 48 h memory RNAi screen.**

The candidate genes are listed along with the used RNAi lines, the 48 h memory performance (PI) and the standard error of the mean (SEM).

**Table S3. Retest of hits with a different RNAi line.**

Results of the 48 h memory experiments, in which hits were retested with a second RNAi line. The tested genes are listed along with the used RNAi lines, the obtained 48 h memory performance (PI) and the corresponding error (SEM).

**Table S4. Predicted *mir-282* target genes.**

List of top 50 *mir-282* target genes predicted by the microRNA.org resource.

